# Global impact of phosphorylation on protein endurance

**DOI:** 10.1101/2020.03.12.989467

**Authors:** Chongde Wu, Qian Ba, Wenxue Li, Barbora Salovska, Pingfu Hou, Torsten Mueller, George Rosenberger, Erli Gao, Yi Di, Yansheng Liu

**Author notes:** These two authors contributed equally to the study (Chongde Wu, Qian Ba).

## Abstract

Post-translational modifications such as phosphorylation can have profound effects on the physicochemical and biological properties of proteins. However, high-throughput and systematic approaches have not yet been developed to assess the effects of specific modification types and sites on protein lifetime, which represents a key parameter for understanding signaling rewiring and drug development. Here we describe a proteomic method, DeltaSILAC, to quantify the impact of site-specific phosphorylation on the endurance of thousands of proteins in live cells. Being configured on the reproducible data-independent acquisition mass spectrometry (DIA-MS), the pulse labeling approach using stable isotope-labeled amino acids in cells (SILAC), together with a novel peptide-level matching strategy, this multiplexed assay revealed the global delaying effect of phosphorylation on protein turnover in growing cancer cells. Further, we identified local sequence and structural features in proximity to the phosphorylated sites that could be associated with protein endurance alterations. We found that phosphorylated sites accelerating protein turnover are functionally selected for cell fitness and evolutionarily conserved. DeltaSILAC provides a generalizable approach for prioritizing the effects of phosphorylation sites on protein lifetime in the context of cell signaling and disease biology, which is highly complementary to existing methods. Finally, DeltaSILAC is widely applicable to diverse post-translational modification types and different cell systems.

## Introduction

Biological signaling features both amplitude and duration^1^. As the prominent molecule for most cellular processes, protein has been characterized by various properties, such as structure, abundance, localization, stability, and turnover. Post-translational modifications (PTMs) generate different proteoforms for a given protein, altering the above properties and leading to diverse functions^2, 3^. Phosphorylation is particular critical PTM and has been shown to be essential for signaling transduction^4^, can mediate protein-protein interaction^5^, and affect the three-dimensional structure of the protein, its thermal stability^6^, and subcellular localization^7^. However, the impact of phosphorylation on protein turnover has so far not been assessed at the proteome-scale.

To adapt to temporal environmental changes, the cells have to utilize kinases and phosphatases to effectively and instantly catalyze phosphate transfer between substrates. Nevertheless, the long-term regulation of endurance and decay of those constitutively phosphorylated proteins is also critical for the cell systems. This regulatory mechanism enables the rewiring of cellular states after adaption^8, 9^, and the establishment of fitness against intrinsic genetic alterations^10, 11^. Specifically, the functional crosstalk between phosphorylation and ubiquitination was discovered to substantially downregulate levels of key phosphoproteins in a variety of pathways such as EGFR/MAPK signaling^9, 12^ and cell-cycle control^13, 14^. However, the high-throughput discovery tools are currently lacking to illustrate how phosphorylated proteins are degraded by proteostasis and proteolysis pathways.

In the last decade, mass spectrometry (MS) based proteomics has greatly facilitated the analysis of proteins and their PTMs^15, 16^. MS-based mapping of phosphorylation and ubiquitylation sites previously suggested that distinct phosphorylation sites often co-occur with ubiquitylation^17^. This analysis, however, relied on the MS detection frequency of PTM sites after individual or tandem PTM enrichments^17^, which does not consider the relative stoichiometry of both PTMs, and the qualitative results are likely affected by the abundance of bulk proteins and the sensitivity of MS analyzers. As an arising MS-based technique, data-independent acquisition mass spectrometry (DIA-MS)^16, 18^ makes the full use of the current high-speed mass spectrometers. DIA-MS usually generates continuous, high-resolution MS2 peak profiles during liquid chromatography (LC) separation that can be used for the simultaneous identification and quantification of thousands of peptides, including their PTMs ^19, 20^. Promisingly, when combined in a pulse experiment using isotope-labeled amino acids in cells (SILAC)^21-24^, we demonstrated that DIA-MS provides high efficiency, reproducibility, and accuracy to measure the turnover rates of proteins^25, 26^ and, recently, their alternative splicing isoforms^27^.

Here we sought to develop an integrative proteomic method that directly interrogates the lifetime of phosphorylated proteins in reference to their unphosphorylated versions. Our method DeltaSILAC (delta determination of turnover rate for modified proteins by SILAC) is built on pulse SILAC (pSILAC) labeling, phosphoproteomic enrichment, and DIA-MS. We show that DeltaSILAC can successfully reveal endurance alterations for constitutively phosphorylated proteins in cellular systems and provides a comprehensive and generalizable approach to characterize the effects of phosphorylation sites on protein lifetime in the context of cell signaling and disease biology.

## Results

### Development of DeltaSILAC for timing the protein endurance in response to site-specific modification

We first overview the rationale of our DeltaSILAC method (**Fig. 1**). In a typical pSILAC experiment, the growing cells are maintained in a steady-state in which the degraded and synthesized protein copies are balanced^28, 29^. Such a concentration balance should also apply to most if not all modified proteins – otherwise, the system will be perturbed. Thus, by culturing cells in the heavy SILAC medium and monitoring the exchange rate of heavy (H) and light (L) amino acids containing peptide signals for a time course, the protein turnover time can be determined by MS analysis for both modified or unmodified proteoforms. Because the concepts of protein lifetime and degradation rate were defined within particular cells and therefore interlace with cell doubling time, herein we postulate the new concept “endurance” for timing the existence of different proteoforms of a given protein and for emphasizing the site-specific effect for a PTM (e.g., a functional phosphorylation) on protein turnover.

**Figure 1.**
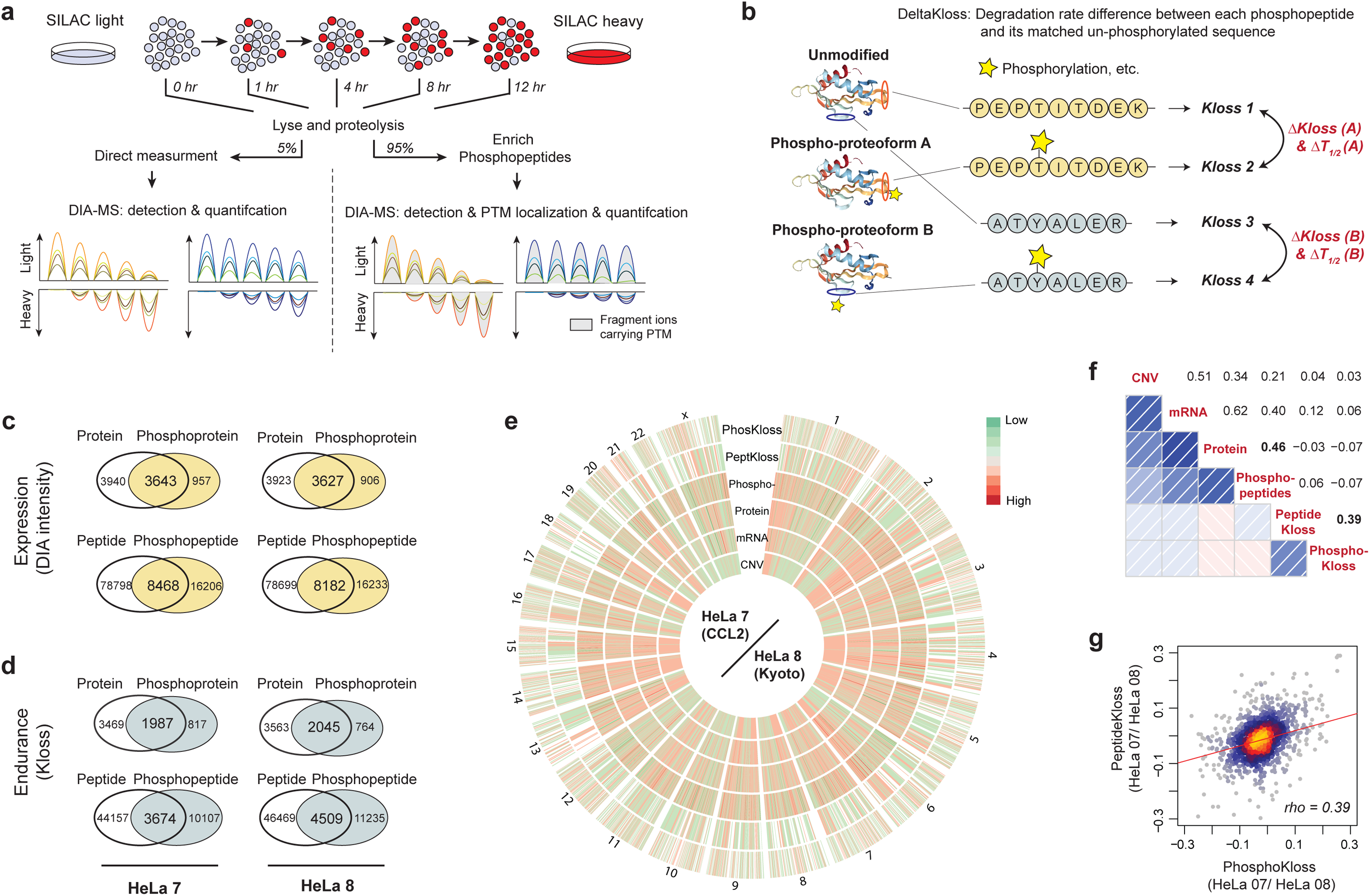
Development of DeltaSILAC to quantify protein endurance with site-specific phosphorylation. **(a)** Experimental workflow illustrating the DeltaSILAC method. The experimental method is comprised of pSILAC labeling, phosphoproteomics, and DIA mass spectrometry (DIA-MS). **(b)** Data analysis strategy illustrating the peptide sequence level matching used in DeltaSILAC. (**c-d**) Venn diagram of overlap between unmodified and phosphorylated proteins and peptides at expression (**c**) and endurance (**d**) levels. **(e)** Circos plot of relative HeLa_7 (CCL2)/ HeLa_8 (Kyoto) ratios in six layers, including expression and endurance, as depicted. Fold changes from high to low are shown in red to green. The data are phosphoproteome centric, i.e., data matched to available phosphoproteomic identifications. **(f)** Spearman correlation analysis between six layers using HeLa_7/ HeLa_8 ratio data. Spearman’s rho is shown, with positive correlations visualized in blue color and negative correlations in red color. **(g)** The scatterplot indicating HeLa_7/ HeLa_8 fold-change ratios of k_Loss_ estimates between matched unmodified and phosphopeptides.

To develop and demonstrate the DeltaSILAC workflow using a well-defined model system, we investigated a panel of 12 HeLa cells initially collected from different research laboratories^26^. A considerable heterogeneity of gene copy number alteration (CNA) was previously documented between these HeLa strains, which leads to systematic rewiring of mRNA, protein, and bulk-protein degradation levels^26^. This prior data provides the basis to assess the roots of phosphoprotein abundance variability. Using DIA-MS^30^, we confidently quantified 24,119 ± 446 Class-I phosphopeptides (with confident phosphorylation site-localization)^4, 31^ across 12 HeLa cell lines (**Fig. S1 & Supplementary Table 1**). Consistent with previous multi-omics reports^32^, we found that phosphoprotein abundance is globally more variable than the corresponding mRNA and protein levels. While we acquired phosphoproteomic profiles for all HeLa CCL2 and Kyoto strains covered by the original study^26^ (**Fig. S1**), we herein chose HeLa_7 and HeLa_8 as a representative HeLa CCL2 and Kyoto strains for the follow-up PTM endurance analysis.

To systematically determine the protein and phosphoprotein lifetime (i.e., endurance in cells), we deployed a five-time point pSILAC experiment at 0, 1, 4, 8, 12 hours for both HeLa_7 and HeLa_8 cells (**Fig. 1**). The cell lysates of different time points were then trypsin proteolyzed. Each peptide aliquot was split, with 5% of peptides used for direct proteomic measurement and 95% for phosphoproteomics. To measure proteomes and phosphoproteomes during labeling, DIA-MS was employed (**Fig. 1a-b**), which provides high quantitative reproducibility for multiple samples labeled by pSILAC^25, 27^. Using a direct identification approach^30^ (see **Methods**), we confidently inferred 7,583 and 7,550 proteins for HeLa_7 and HeLa_8 from single-shot measurements, detected by 87,266 and 86,881 peptides (both peptide- and protein-FDR were < 1%). For 5,456 and 5,608 proteins (detected by 47,831 and 50,978 peptides), we computed turnover rates (**k**_**Loss**_), which can be mathematically transformed to provide an estimation of the protein half-life time (T_50%_ values^33^, hereafter, **T**_**1/2**_, **see Methods and Fig.1c-d & Fig. S2**). Moreover, among ∼24,000 phosphopeptides detected in HeLa_7 and HeLa_8, 13,781 and 15,777 phosphopeptides (that is, 12,134 and 13,078 phosphosites) were quantified with a T_1/2_. We found that T_1/2_ correlation between HeLa_7 and HeLa_8 for phosphopeptides (R=0.90) was similar as T_1/2_ for proteins (R=0.86, **Fig. S2**), suggesting the quantitative performance of pSILAC-DIA^25, 27^ can be extended to phosphoproteomics. Altogether, this dataset **(Supplementary Table 2)** presents a first systematic analysis of phosphoproteome endurance.

We then benchmarked the phosphopeptide dynamics in comparison to other omics layers. For this purpose, the genome-wide ratios between HeLa_7 and HeLa_8 were assessed across the levels of gene copies, mRNA, bulk-protein expression, phosphopeptides, as well as turnover rates of all peptides and phosphopeptides (**Fig. 1e**). We found that whereas the change of phosphopeptide abundance follows mRNA and protein levels (R=0.40 and 0.46), the turnover regulation of phosphopeptide does not (**Fig. 1f**). Additionally, the correlation between turnover ratios for phosphopeptides and peptide counterparts are modest (R=0.39; **Fig. 1g**), similar to that correlation between abundance ratios (R=0.46). This data, therefore, highlights the strong need to independently analyze phosphopeptides for both abundance and endurance. Also, although the gene dosage compensation mechanism at the protein turnover level was confirmed by positive mRNA∼peptide k_Loss_ correlation (R=0.12), we found the mRNA∼phosphopeptide k_Loss_ correlation to be weaker (R=0.06) which is not significant after correction against their relevance to peptide k_Loss_ (p>0.05). This implicates that broad CNA have a limited impact on phosphoprotein endurance overall. We therefore analyzed on HeLa_7 and HeLa_8 datasets independently in the following analysis.

To summarize, with DeltaSILAC, we were able to measure half-lives for both modified and unmodified versions of the same protein (**Fig. 1d**). This matched dataset enables a novel strategy (see below) to interrogate the impact of PTM on protein expression duration.

### Regulation of protein endurance by phosphorylation is site-specific, rather than gene-specific

To pinpoint PTM influence on protein lifetime, we subtracted T_1/2_ of backbone sequence-matching non-phosphorylated peptides (**np-peptide**) from the T_1/2_ of counterpart phosphopeptides (**p-peptide**), resulting in the delta value (ΔT_1/2_). Thus, ΔT_1/2_ was used for all following endurance analysis. After filtering (**see Methods)**, 1,919 and 2,147 such pairs of np- and p-peptides in HeLa_7 and HeLa_8 were quantified. We found that for those proteins identified with multiple phosphorylation sites, different sites can lead to a variable ΔT_1/2_ (**Fig. 2**). For example, the pT490 increased T_1/2_ of AHANK by ∼7.5 hours (i.e., ΔT_1/2_ = 7.71 and 7.58 hours in HeLa_7 and HeLa_8) whereas pT4100 shortened T_1/2_ by > 15 hours (i.e., ΔT_1/2_ = −15.61 and −16.86 hours; **Fig. 2a**). For MAP4 certain phosphosites such as pS507 and pT521 could greatly enlarge T_1/2_ by > 40 hours, having a much higher protein endurance modulation effect than other phosphosites of the same protein (**Fig. 2b**). In comparison, for MARCKS, phosphorylation on most sites detected on average similarly increased protein endurance of about 5 hours (**Fig. 2c**). Interestingly, all the 19 phosphorylated versions of SF3B1 barely changed their lifetime as compared to the non-phosphorylated counterpart (ΔT_1/2_ =0.7194 ± 1.956 hours) and other cases (**Fig. 2d-e**), indicating endurance can be robust for specific proteins despite variable modifications.

**Figure 2.**
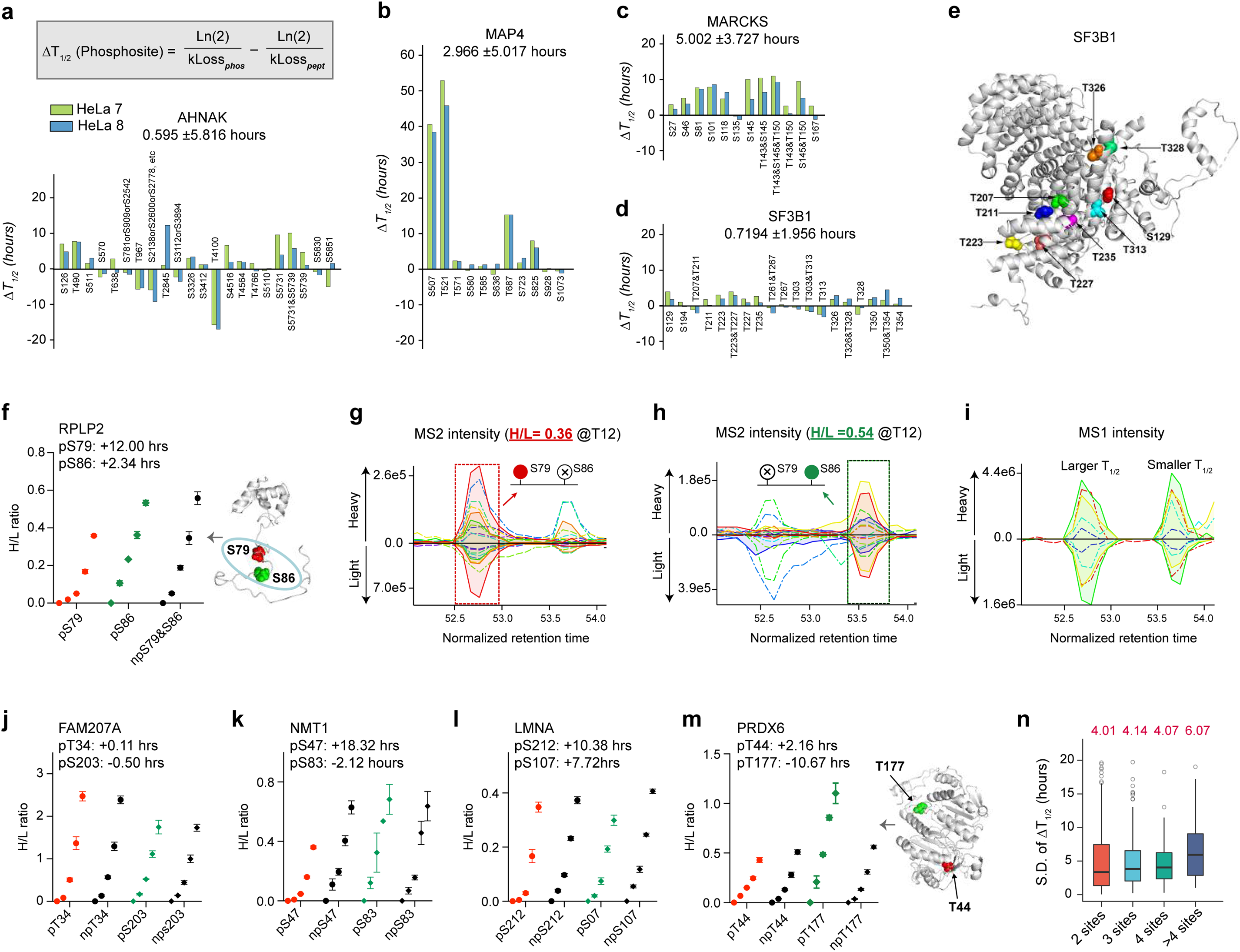
Regulation of protein endurance by phosphorylation in DeltaSILAC data. (**a-d**) The averaged ΔT_1/2_ values (N=3 biological replicates) determined by an independent analysis of HeLa_7 and HeLa_8 cells for protein examples of multiple phosphorylation sites. These examples include **(a)** neuroblast differentiation-associated protein AHNAK (AHNAK), **(b)** Microtubule-associated protein 4 (MAP4), **(c)** Myristoylated alanine-rich C-kinase substrate (MARCKS), and **(d)** Splicing factor 3B subunit 1 (SF3B1). **(e)** Demonstration of localization of nine phosphorylated sites of SF3B1 in its known structure. **(f)** The Heavy-to-Light (H/L) ratios determined for S79 and S86 phosphorylated form (p-peptide) and their shared un-phosphorylated form (np-peptide, LASVPAGGAVAVSAAPGS_79_AAPAAGS_86_APAAAEEK) in 60S acidic ribosomal protein P2 (RPLP2). **(g)** The DIA MS/MS peak groups during LC elution containing unique fragment ions for the peptide carrying phosphorylated S79 (pS79). Note that the heavy (above middle zero lines) to light peaks (below zero line) are scaled, respectively. **(h)** The DIA MS/MS peak groups during LC elution containing unique fragment ions for the peptide carrying phosphorylated S86 (pS86). Note that the heavy to light peaks are scaled, respectively. **(i)** The MS1 peak groups from the same DIA run during the same elution region (corresponding to a peptide containing single phosphorylation). **(j-m)** Individual examples of H/L ratios determined for paired p-peptide and np-peptide for **(j)** Protein FAM207A, **(k)** Glycylpeptide N-tetradecanoyltransferase 1 (NMT1), **(l)** Prelamin-A/C (LMNA), and **(m)** Peroxiredoxin-6 (PRDX6). **(n)** Data variability of ΔT_1/2_ values summarized as standard deviation (S.D.) for different phosphosites within the same protein. All proteins were classified by the number of phosphosites measured by DeltaSILAC. The red numbers denote the median values in each group.

We further found that our approach was able to impressively discriminate the turnover discrepancy between proteoforms carrying closely located phosphosites, even those identified on the same peptide backbone. The sites S79 and S86 in RPLP2 present such an example, as they share the same np-peptide **(Fig. 2f)**. The unique MS2 signatures carrying either pS79 or pS86 were extracted from the DIA-MS dataset. Moreover, the heavy-to-light (H/L) ratios of the two sites during pSILAC labeling were confidently resolved to respective LC peak groups, which have only a < 1 min retention time (RT) interval that presents a challenging case for MS1 alignment^34, 35^ (**Fig. 2g-h**). Accordingly, a T_1/2_ difference of 9.66 hours between pS79 and pS86 was successfully determined. In contrast, we found that the MS1 features alone cannot differentiate the two phosphopeptides owning to their identical m/z **(Fig. 2i)**.

To gauge the precision of ΔT_1/2_, we checked the H/L ratios of np-peptides and p-peptides during pSILAC labeling. We found the sequence-matching strategy to be useful, because even np-peptides can show variable labeling rates, such as npT34 and npS203 of FAM207A in **Fig. 2j**. The extended protein endurance by phosphosites was credibly inferred from H/L ratios measured by DIA-MS, such as pS47 of NMT1 and pS212 & pS107 of LMNA (**Fig. 2k-l**). Other sites instead reduce the proteoform endurance, such as pT177 in PRDX6 (**Fig. 2m**). We further found the determined np-peptide and p-peptide T_1/2_ to be highly stable across replicates, validating the reproducibility of experimental and computational pipeline in DeltaSILAC and indicating that even small changes in protein endurance (for example, ∼1 hour) are detectable with statistical cutoffs (**Fig. S3**). Therefore, DeltaSILAC reliably uncovered a 4-6 hours variability of half-life time, on average, between differentially phosphorylated versions of the same protein (**Fig. 2n**).

Altogether, our analysis demonstrates that DeltaSILAC can accurately assess the effects of phosphorylation on protein endurance in a site-specific manner.

### Phosphorylation prefers to increase endurance for many proteins in growing cells

To understand the global effect of phosphorylation on protein turnover, we analyzed T_1/2_ in three comparative scenarios. **1.** Total comparison: T_1/2_ of all phosphopeptides identified *versus* T_1/2_ of all bulk proteins identified **(Fig. 3a-b, left panel)**. This “unmatching” comparison is regardless of the mapping of np- or p-peptides. **2.** Protein-matched comparison: T_1/2_ of phosphopeptides whose corresponding proteins were identified *versus* T_1/2_ of proteins whose corresponding phosphosites were identified **(Fig. 3a-b, middle panel)**. This comparison thus excludes ∼10,000 phosphopeptides and ∼3,500 proteins that were only detected by either phosphoproteomics or proteomics (**Fig. 1d**). **3.** Peptide-matched comparison: T_1/2_ of p-peptides *versus* T_1/2_ of np-peptides **(Fig. 3a-b, right panel)**. This comparison only accepts T_1/2_ data with a pair of np- and p-peptides. Comparison 1 suggests that the total phosphoproteome and the proteome have comparable T_1/2_ (median is 16.8 vs. 16.3 hours for HeLa_7 and 15.8 vs. 16.1 hours for HeLa_8). The slightly reduced T_1/2_ in HeLa_8 from HeLa_7 is expected because HeLa Kyoto has a shorter doubling time than HeLa CCL2, as reported^26^. Intriguingly and significantly, a trend of increasing protein endurance conferred by phosphorylation was demonstrated by both protein- and peptide-matched strategies: the T_1/2_ median increase by phosphorylation was determined to be 0.70 and 1.10 hours for HeLa_7 and HeLa_8 according to the protein-matched comparison and 2.53-2.55 hours according to peptide-matched comparison (Comparison 2 and 3; **Fig. 3a-b**). The volcano plots suggest that, based on the matching-peptide comparison, T_1/2_ is significantly upregulated for 623 and 679 phosphopeptides but only downregulated for 148 and 156 in two cell lines (P < 0.01, T_1/2_ change > 1.5 hours; **Fig. 3c-d**). The different outcome from Comparison 1 might be ascribed to the phosphoproteomic enrichment step. This step could cover more low-abundant proteins, which in turn have a shorter T_1/2_ than high-abundant proteins detected by total proteomics^25, 29, 36^ (**Fig. S4a**). The above global result further emphasizes the importance of using sequence-matched controls for analyzing the endurance of modified proteoforms.

**Figure 3.**
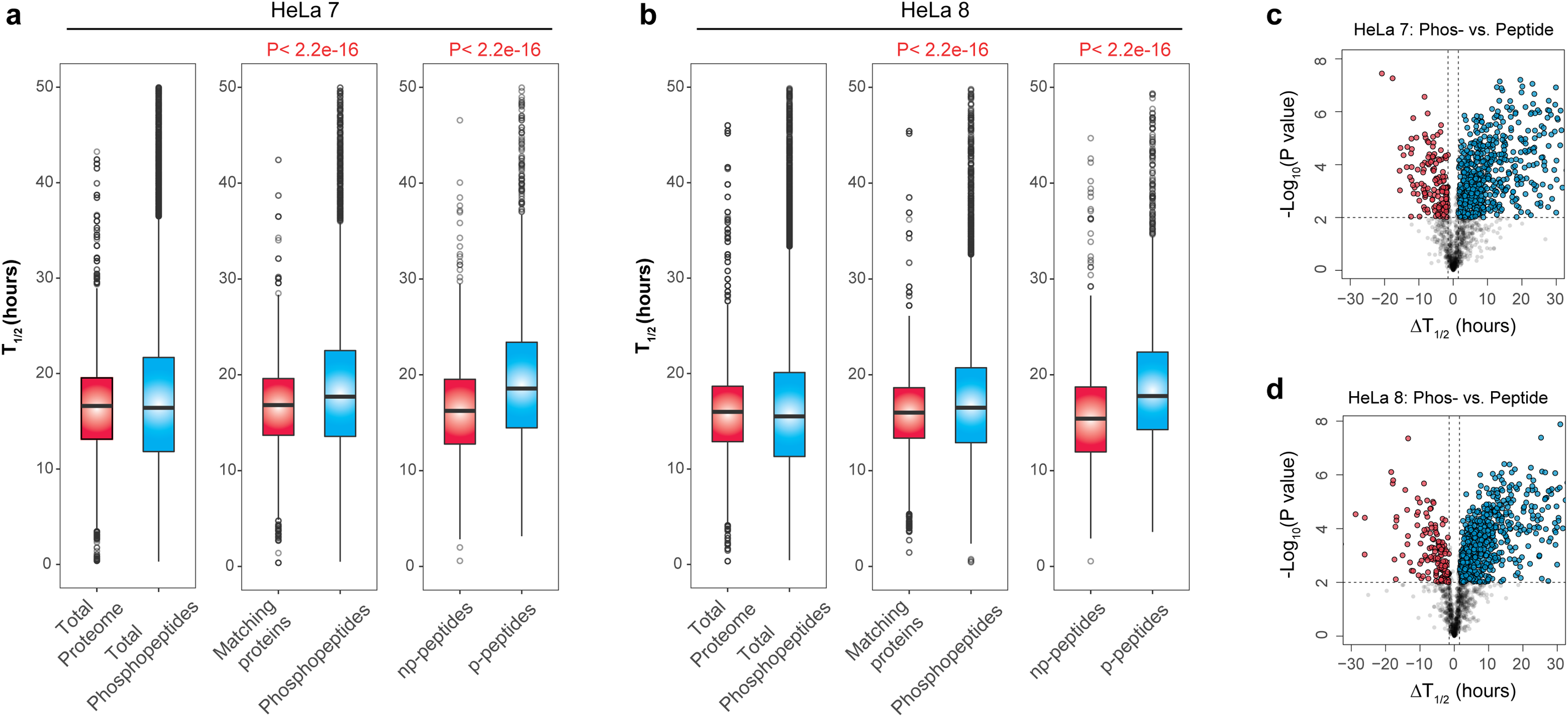
Phosphorylation increases endurance for the majority of proteins in growing HeLa_7 and HeLa_8 cells. (**a-b**) Distribution of T_1/2_ values in total proteome *versus* phosphopeptides matched proteins *versus* phosphoproteins (based on individual phosphopeptides not summarized), and matched peptides *versus* phosphopeptides in HeLa_7 (**a**) and HeLa_8 (**b**) cells. *P* values were calculated by Wilcoxon sum test. The borders of the box represent the 25th and 75th percentile, the bar within the box represents the median, and whiskers represent the range. (**c-d**) Volcano plots for ΔT_1/2_ of 3 vs. 3 biological replicates in HeLa_7 (**c**) and HeLa_8 (**d**) cells. P values were calculated by Student’s T-tests. The color dots denote the significantly altered gene expressions (P < 0.01; ΔT_1/2_ > 1.5 hours).

Taken together, the matching comparisons somewhat surprisingly elaborate that many proteins tend to increase rather than decrease T_1/2_ when phosphorylated. The peptide-matching strategy (Comparison 3) which generated more significant ΔT_1/2_ values is thus preferably used, whereas the protein-matching strategy (Comparison 2) was also adopted in some of the following analyses because it generates 2.5 times more the valid ΔT_1/2_ values for phosphosites (6,834 for HeLa_7 and 7,654 for HeLa_8).

### Structural features associated with protein endurance altered by site-specific phosphorylation

The effects of phosphosites on protein endurance (**Fig. S5a**) prompted us to investigate whether certain local structural environments and other properties of particular phosphosites are associated with ΔT_1/2_. By comparing site-specific T_1/2_ to a recently published dataset reporting the site-specific melting temperature (T_m_) for phosphoproteins^6^, we found a positive correlation of R=0.20 across all phosphosites. This relationship holds even after stringent correction using bulk-protein abundance which positively correlates to both T_1/2_ and T_m_ (**Fig. S4)**. Conceivably, this result indicates that phosphorylation sites bringing more thermal stability to proteins may increase protein endurance (**Fig. 4a**) and might be suggestive of a coupling mechanism between structural stability and expression stability for achieving cellular proteostasis.

**Figure 4.**
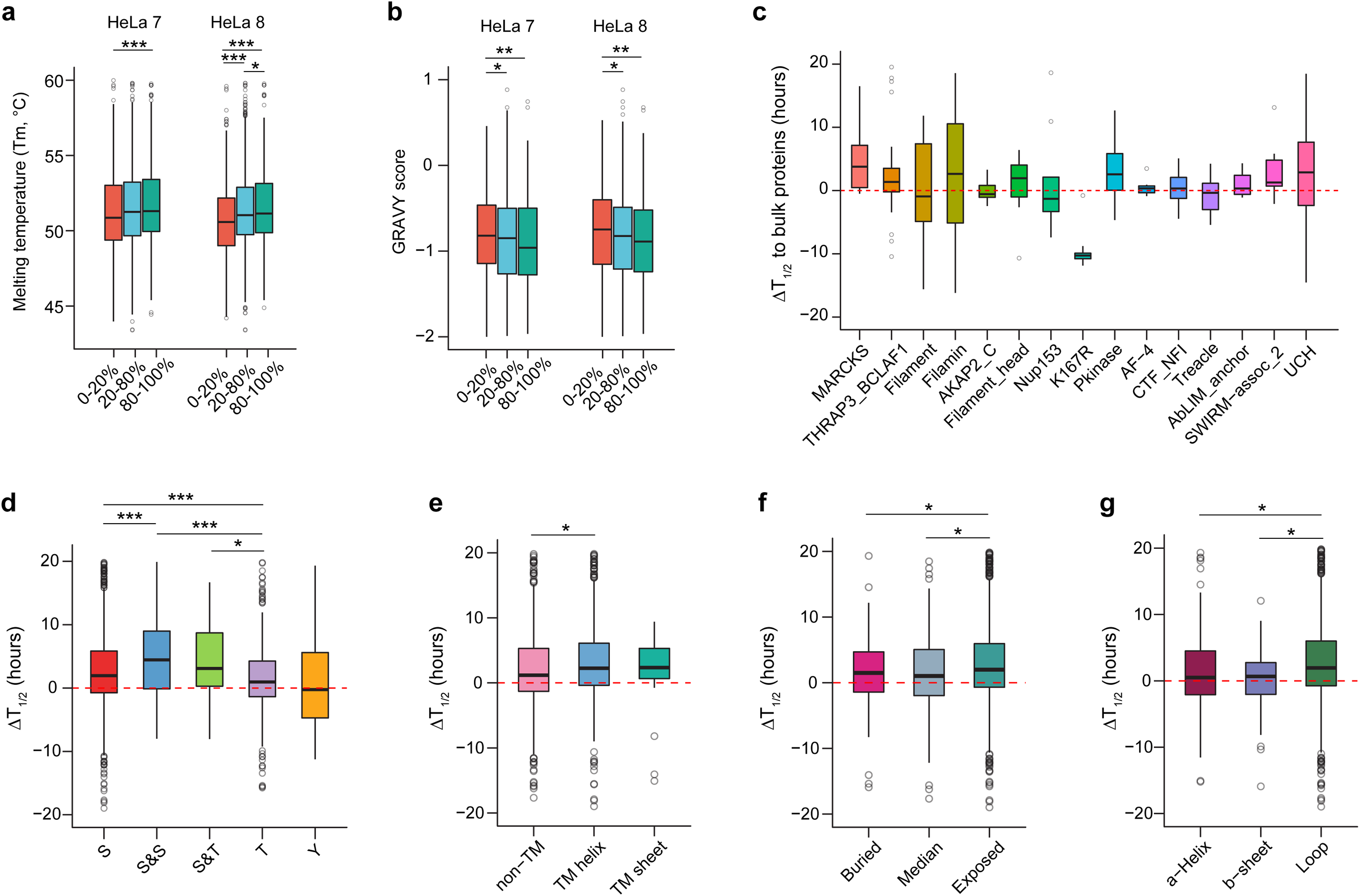
Global relationships between local structural features and phosphorylation-altered protein endurance. **(a)** The melting temperature (T_m_, °C) of matched phosphosite-specific proteoforms is distributed into T_1/2_ quintiles from small to large in HeLa_7 and HeLa_8 cells (***P < 0.001, *P < 0.05, Wilcoxon sum test). **(b)** The GRAVY score of phosphopeptides (based on their naked sequences) is distributed into ΔT_1/2_ quintiles from small to large percentage in HeLa_7 and HeLa_8 cells (**P < 0.01, *P < 0.05, Wilcoxon sum test). **(c)** Comparisons between the most detected protein domains and ΔT_1/2_ values (to bulk proteins) for measured phosphosites in HeLa_7 cells. **(d)** Distribution of ΔT_1/2_ values for phosphopeptides with indicated phospho-amino acids and combinations thereof in HeLa_7 cells (***P < 0.001, *P < 0.01, Wilcoxon sum test). (**e-g**) Comparisons of ΔT_1/2_ values and predict transmembrane topologies (**e**), solvent accessibility (**f**), and predicted secondary structure elements (**g**) in HeLa_7 cells (*P < 0.05, Wilcoxon sum test). All corresponding results in HeLa_8 can be found in Figure S6-7.

Next, to facilitate the relative ΔT_1/2_ analysis, we divided ΔT_1/2_ into quintile segments, with Q1 (0-20%) representing phosphosites reducing endurance (i.e., faster turnover), Q2-Q4 (20-80%) representing intermediately regulated cases, and Q5 (80-100%) representing the highest endurance (i.e., dramatically slowed down protein turnover with phosphorylation; **Fig. S5b-d**). *Firstly*, we found that the grand average of hydropathy (GRAVY) score of all np-peptides negatively correlates to ΔT_1/2,_ suggesting the higher peptide-hydrophobicity tends to stabilize the existence of the phospho-proteoform (*P*= 0.0029 and 0.0014 between Q1 and Q5 in HeLa_7 and HeLa_8, Wilcoxon sum test; **Fig. 4b**). *Secondly*, different protein domains did not show a general effect on ΔT_1/2_ (**Fig. 4c & Fig. S6**). One of the exceptions is the increased endurance for phosphorylated MARCKS (*P*= 0.006 and 0.034), an intrinsically disordered, alanine-rich protein whose phosphorylation translocates the protein from plasma membrane to cytoplasm through conformational changes^37^. Another exception is the shortening endurance of phosphorylated KI67/Chmadrin repeat (*P*= 4.3E-09 and 1.5e-4) with presumable functions in the cell cycle process^38^. *Thirdly*, global ΔT_1/2_ values among the phosphorylated serine (S), threonine (T), and tyrosine (Y) sites revealed longer lifetime for pS than pT containing peptides (median is 2.43 vs. 1.48 hours of ΔT_1/2_, *P*=0.0011, for HeLa_7). Seemingly, pY yields the shortest endurance (ΔT_1/2_ = 0.066 hours). Notably, the combination of two phosphorylated amino acids further increased protein endurance: ΔT_1/2_ is 6.87 hours for two pS containing peptides and 3.70 for those with one pS and one pT (**Fig. 4d**), higher than single phosphosite containing peptides. This reinforces the delaying effect of individual phosphorylation on protein turnover globally. All identical observations were obtained from HeLa_8 (**Fig. S7a**). *Finally*, phosphosites without transmembrane topologies (non-TM) demonstrated smaller ΔT_1/2_ than those with transmembrane domains (e.g., helices, **Fig. 4e, and Fig. S7b**). Nevertheless, phosphosites predicted in a buried region with low solvent accessibility and in helical, or β-sheet demonstrated smaller half-life changes than those in exposed and loop area, in agreement to thermal stability data^6^ (**Fig. 4f-g and Fig. S7c-d**).

In summary, we discovered that local protein structural properties around specific phosphorylation sites influence protein endurance in response to phosphorylation.

### Functional insights for altered protein endurance induced by phosphorylation

Next, we studied the functional relevance between phosphorylation and protein endurance alteration. Ochoa et al. recently conducted a meta-analysis of phosphoproteomic datasets and extracted 59 features annotating phosphosite functions. They further integrated these features to a single score that prioritizes phosphosites relevant for cell fitness^11^. Correlating ΔT_1/2_ to this score reveals a strong, negative trend (**Fig. 5a**), reassuringly indicating that those functionally important phosphosites are likely to have a faster turnover, underscoring the actual functional relevance of previous studies focusing on the crosstalk between phosphorylation and ubiquitination as well as phosphodegrons^17, 39^. In-depth correlation analysis suggests that ΔT_1/2_ has high evolutionary relevance. For example, compared to Q2-4 and Q1, Q5 sites exhibit a much higher variant tolerance (SIFT score) that summarizes the lower conservation for phosphosite residue to alanine mutations or any mutations^11, 40^ (**Fig. 5b & Fig. S8**). Thus, phosphosites, mainly slowing down protein turnover, tend to be less conservative during evolution. Moreover, the kinase and kinase-substrate annotation suggested that phosphorylated kinases generally increased their lifetime, just as other phosphorylated proteins. Deviating from this norm are a few examples showing faster turnover in steady-state cells, such as Rho-associated coiled-coil containing kinases, ROCK1, and ROCK2 (**Fig. 5c &S9**). Kinase-substrate mapping further revealed that the phosphosites activated by Cyclin-dependent kinases, especially CDK1, have a drastically short endurance (**Fig.5d & Fig. S10a**), in agreement with prior knowledge that the phosphorylation coupling degradation is essential in cell division^39^. In the next step, to obtain an unbiased functional view for site-specific ΔT_1/2_, we performed a biological process (BP) enrichment analysis in each segment from Q1 to Q5 (**Fig. 5e** for HeLa_7 **& Fig. S11** for both cells). A few BPs are universally enriched by the phosphoproteome (throughout Q1-Q5), such as cell-cell adhesion, RNA splicing, and RNA binding. Specific BPs essential for translational control such as translational initiation (P=0.035), protein biosynthesis (P=0.013), as well as cell cycle (P=0.008) are enriched in Q1, demonstrating their shorter endurance upon phosphorylation. The SH3 domain was preferably enriched in Q2-4, indicating the robustness of endurance for these phosphoproteins, on average. Q5 enriched distinctive BPs such as translocation (P=1.93e-5), glucose transport (P=4.1e-4), and tRNA transport from the nucleus (P=6e-4), suggesting a stabilization of the expression of proteins participating in cellular transport by phosphorylation. In addition, distributing ΔT_1/2_ values to their protein subcellular locations^41^ only revealed an exceptional case of Nuclear Speckles, where the phosphoproteins have a relatively faster turnover (**Fig. S12**).

**Figure 5.**
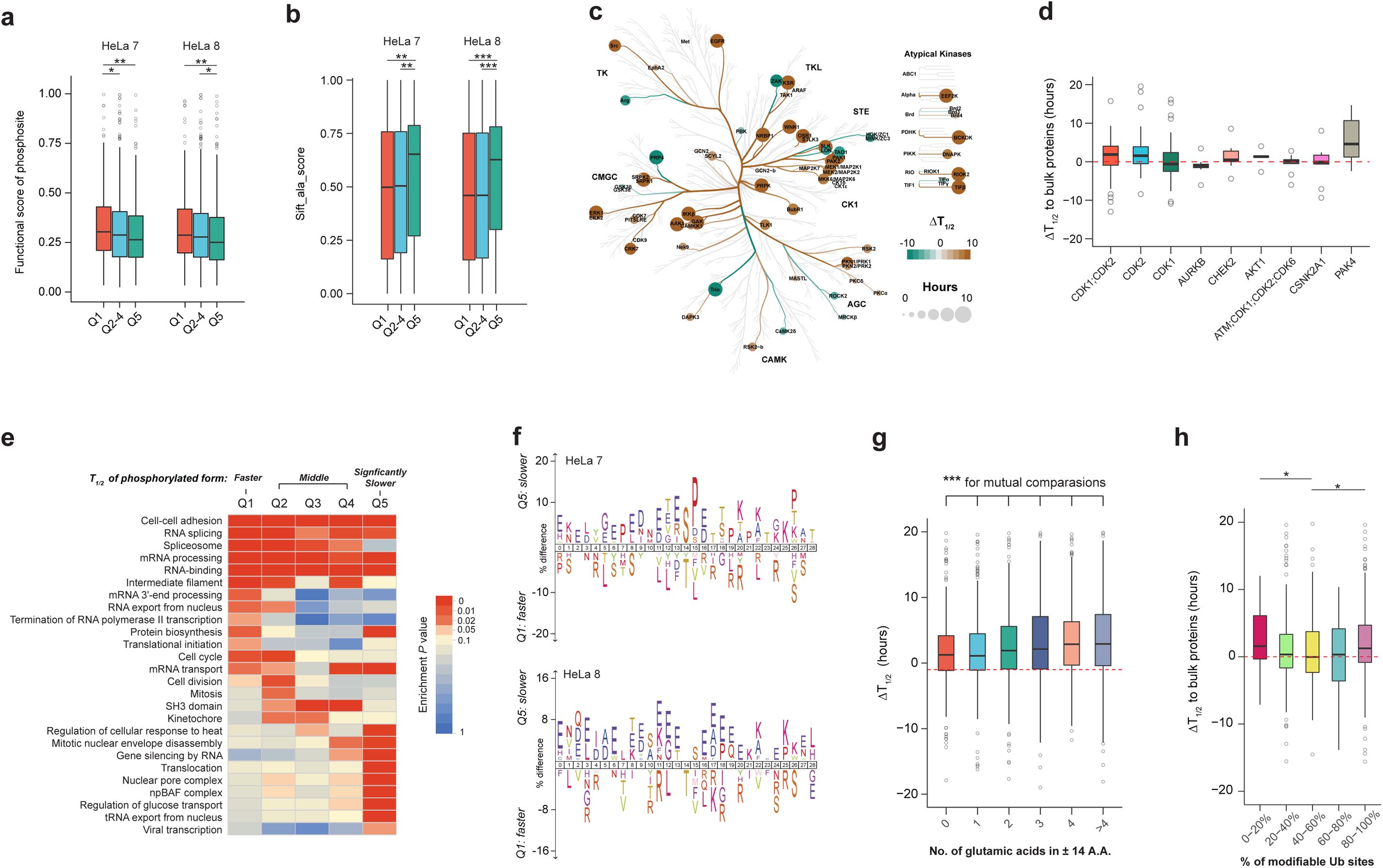
Functional features associated with phosphorylation-altered protein endurance. **(a)** The functional scores of phosphosites are distributed into ΔT_1/2_ quintiles (Q1 to Q5: small to large) in HeLa_7 and HeLa_8 cells (**P < 0.01, *P < 0.05, Wilcoxon sum test). **(b)** The Sift scores of the alanine mutant for phosphosites are distributed into ΔT_1/2_ quintiles (Q1-Q5: from small to large) in HeLa_7 and HeLa_8 cells (***P < 0.001, **P < 0.01, Wilcoxon sum test). **(c)** Depiction of human kinome indicating all kinases that harbor phosphorylation site with less (green) or greater (brown) phosphoproteoform T_1/2_ value compared to bulk proteins in HeLa_7 cells. A large node represents a greater difference. For kinase with multiple phosphosites, only the largest change (maximum absolute ΔT_1/2_ value) was shown. **(d)** Comparisons between detected kinase substrate motifs and ΔT_1/2_ values (to bulk proteins) for identified phosphosites in HeLa_7 cells. **(e)** Functional processes enrichment analysis for all five quintiles of proteins according to the ranked ΔT_1/2_ value of phosphoproteoforms compared to the matched proteins in HeLa_7 cells. **(f)** Sequence analysis of the ±14 amino acids around the phosphosite with the smallest and longest ΔT_1/2_ values (Q1 vs. Q5, where Q1 has a faster turnover for p-peptides than np-peptides, and Q5 has a much slower turnover for p-peptides than np-peptides) in HeLa_7 and HeLa_8 cells. The percentage of significant residues (P < 0.05) were shown. **(g)** Distribution of ΔT_1/2_ values of phosphopeptides with different numbers of Glutamic acids (Es) around the phosphosite (±14 amino acids) in HeLa_7 cells (***P < 0.001, Wilcoxon sum test). **(h)** Distribution of ΔT_1/2_ values (to bulk proteins) with different percentages of modifiable ubiquitination sites (the number of ubiquitination sites over the total number of Lys) around the phosphosite (±14 amino acids) in HeLa_7 cells (*P < 0.05, one-sided Wilcoxon sum test). All corresponding results in HeLa_8 can be found in Figure S8-11.

Besides biological function annotation, we assessed the frequency of amino acids (A.A.) surrounding phosphosites, with the expectation to identify more coupling rules for phosphorylation and degradation (**Fig. 5f**). The central A.A. position in this analysis confirmed that pT and pY outperform pS in reducing protein endurance (see also **Fig. 4d**). Also, interestingly, we did not detect apparent enrichment of lysine (K) residues around phosphosites in Q1, although lysine residues can be potentially ubiquitylated. Instead, we discovered a prevalent enrichment of glutamic acid (E) in Q5. Indeed, more Es within ±14 A.A. residues remarkably increase the phosphosite expression stability, which was not previously reported (**Fig. 5g and Fig. S10b**). Finally, we mapped the phosphosites with their ΔT_1/2_ to a dataset identifying ubiquitylation peptides containing K-ε-diglycine^42^ in human cells (diGly, **Fig. 5h and Fig. S10c**). We found that ΔT_1/2_ indeed shrinks with an increasing percentage (from 0-60%) of modifiable Ks^17, 43^ around the phosphosites. However, the highest density of K(diGly) conversely confers delayed degradation effect (i.e., ΔT_1/2_ is larger for 80-100% range, **Fig. 5h**), suggesting that the density of K(diGly) alone without the ubiquitin stoichiometry is not enough to denote protein degradation extent.

In summary, biological annotation on DeltaSILAC data provides the novel functional recognition of phosphorylation stability.

### Regulation of protein endurance due to phosphorylation can be largely conserved across cells

Above, we analyzed cervical cancer cells HeLa_7 and HeLa_8 by DeltaSILAC. We next sought to assess the generalizability of the major observations in two colorectal cancer (CRC) cell lines, SW948 and RKO, by conducting DeltaSILAC measurement. We performed absolute and relative correlation analysis for T_1/2_ and ΔT_1/2_ between all four cell lines (**Fig. 6a-c**). The correlations of proteome-wide, as well as phosphoproteome-wide T_1/2_ between two CRC cells and two HeLa strains, were higher than correlations across the tissue types of the cells (R=0.82-0.83 for CRC cells, R=0.87-0.89 for HeLa cells, in contrast to R=0.53-0.59 between them). The relative measure, ΔT_1/2_, resulted in almost comparable correlations (R=0.76 for CRC cells, R=0.83 for HeLa cells, and R=0.31-0.49 between them). Also, we were able to broadly reproduce observations made in HeLa cells, such as the enrichment of Es in Q5 (**Fig. 6d**) and the globally increased protein endurance by phosphorylation (**Fig. S13**). Thus, ΔT_1/2_ demonstrates a promising, non-stochastic parameter to be measured for different cell lines.

**Figure 6.**
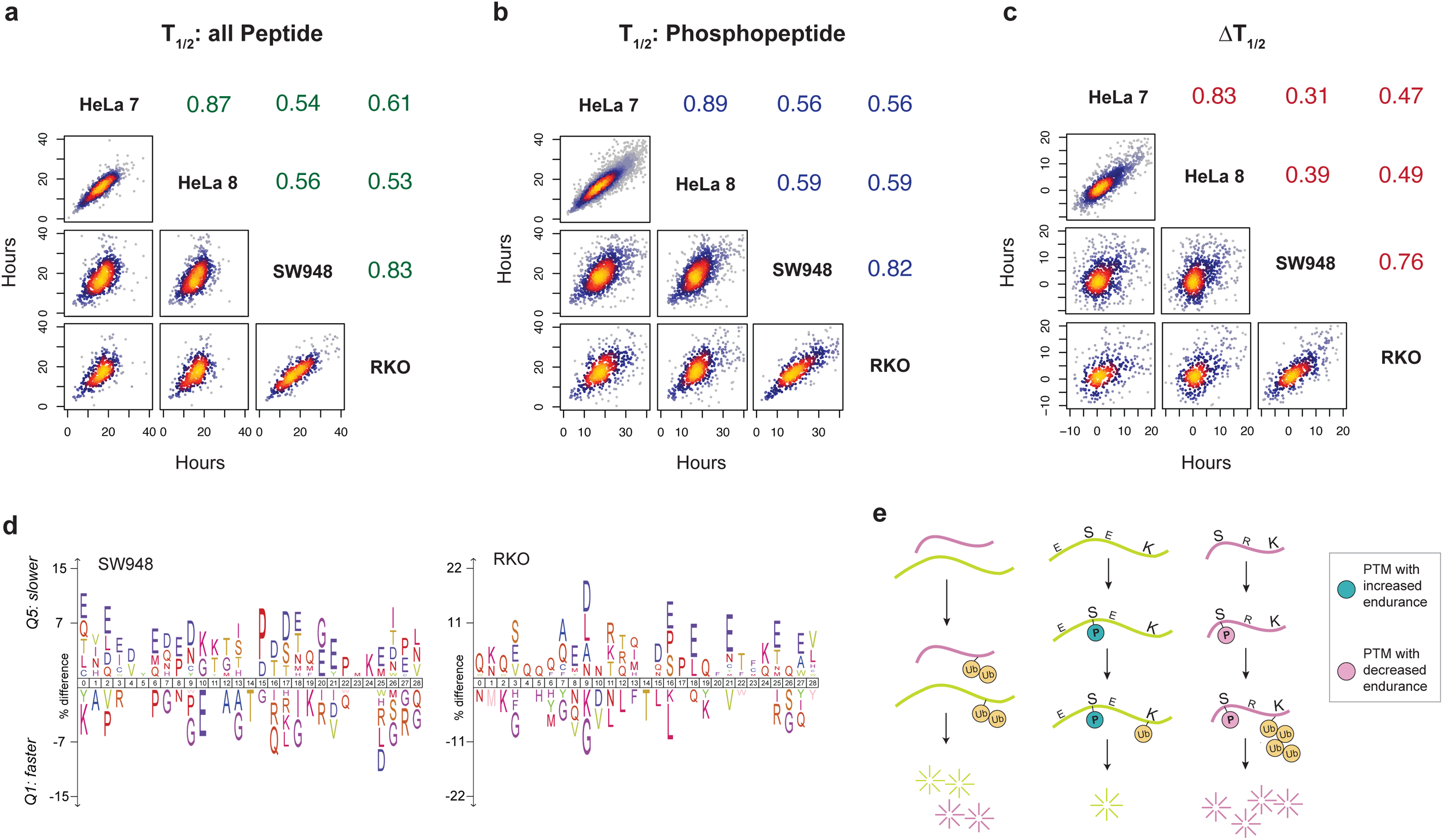
Phosphorylation conferred protein endurance regulation across HeLa_7, HeLa_8, SW480, and RKO cells. (**a-c**) Spearman’s correlation among four cell lines of T_1/2_ values for all peptides (**a**), phosphopeptides (**b**), and ΔT_1/2_ values (**c**). **(d)** Sequence analysis of the ±14 amino acids around the phosphosite with the fastest and slowest ΔT_1/2_ values (Q1 vs. Q5) in SW948 and RKO cells. The percentage of significant residues (P < 0.05) were shown. **(e)** Proposed model coupling protein endurance to phosphorylation with different surroundings.

## Discussion

Here we established an unbiased proteomic method, DeltaSILAC, that quantitatively measures the site-specific impact on protein turnover rendered by PTM events on thousands of proteins. Although phosphorylation was heavily studied in response to perturbations such as drug treatment and was heavily discussed together with instant “phosphate-transfer” activities by kinase and phosphatase, many druggable phosphorylation sites are constitutive, stabilized, or rewired in the disease status. We used the fact that in steady-state growing cells, the abundance of almost all proteins, as well as their modified versions, achieves a balance between synthesis and degradation^22, 24, 29^. Accordingly, a pSILAC experiment provides an excellent opportunity to study the *de facto* outcome of this balance for proteins and proteoforms. DeltaSILAC essentially measures this balance *after* the addition and removal of phosphate groups by kinases and phosphatases and the stabilization of the protein synthesis and degradation processes such as the ubiquitin-proteasome pathway and lysosomal proteolysis. This is because, in DeltaSILAC, the counterpart non-modified peptide (np-peptide, or at least the deriving protein) for each Class-I phosphopeptide has to be quantified with a half-life, simulating a virtual, non-modified proteoform reference that is also PTM site-specific. It should be stressed, thus, that the *bona fide* endurance change at the protein level might be smaller than ΔT_1/2_ appeared, due to the mutually causal relationship between lifetimes of modified and non-modified proteoforms in growing cells. The real proteoform endurance, in theory, has to be measured by top-down approaches^3^, which currently still lack sensitivity. Nevertheless, we believe DeltaSILAC presents a significant methodological advance, in contrast to traditional qualitative LC-MS/MS studies^17^ that might be biased by PTM enrichment efficiency, basic protein abundance, and the possibility of other co-existing PTMs.

Of note, the high reproducibility of DIA-MS is extremely important for DeltaSILAC, because DIA-MS favorably supports the pSILAC experiment, which routinely requires multiple samples to be measured. The single-shot DIA-MS already achieved substantial coverage on both proteome and phosphoproteome, minimizing the overall instrument time needed for a global analysis. Importantly, as illustrated in **Fig. 2f-i**, the MS2-level ion signatures unique to PTM can be extracted from DIA, together with high-resolution LC separation, to reach unprecedented analytical specificity for each PTM sites.

Such specificity may be challenging to achieve for MS1 alignment algorithms^34, 35^. Another prominent advantage of DeltaSILAC is its accuracy because every H/L ratio is inferred from the same MS2 scans in the same MS injection. This means, even MS has a sensitivity drift between injections, the relative H/L ratios between PTM and non-PTM peptides, once identified, remain quantitatively accurate.

We show that DeltaSILAC has revealed novel biological insights. *Firstly*, the majority of the phosphorylation sites were determined to increase their lifetime, at least in the four investigated cancer cells under stable growing status. This delayed turnover of phosphoproteins, in the first sight, seems to be contradictory to the functional crosstalk events discovered between phosphorylation and ubiquitylation^8, 9, 12^. The further scrutiny suggests that the phosphosites of faster turnover (i.e., those in Q1) are indeed functionally more essential for cellular fitness, more evolutionarily conserved, and enriched for CDK1 substrate sites, all consistent to previous findings^8, 9, 13, 14^. Thus, the discovery that phosphorylation often negatively acts on protein turnover might be an observation that was underrepresented in previous studies due to the lack of unbiased quantitative measurements. *Secondly*, local protein structural feature analysis and biological annotation illustrate that these turnover-delaying phosphoproteins may involve multiple-phosphorylated sites, frequently in the loop and exposed regions, and might significantly regulate cellular transport and various processes. For example, the nuclear pore complexes (NPC) were previously identified to maintain over a cell’s life through slow but finite exchange^44^. With a few sites being phosphorylated, these NPC proteins were found by DeltaSILAC to resist degradation even more, reinforcing their roles in cellular ageing^44^. *Thirdly*, the dataset suggests the lack of glutamic acid residues around the phosphosite might provide a more prominent feature than the existence of lysine in predicting the phospho-proteoform endurance. The frequency of glutamic acid is remarkably linked to longer phosphosite endurance. The above observations, although preliminary, open an attractive avenue for perturbating the activity of key phosphorylation sites that are known to be responsible for drug response and disease development, for the possibility of phenotype management (**Fig. S14**). The targeted management could be directed by the prediction rules^39^ **(Fig. 6f)** exemplified above. Generating a data inventory of endurance features by DeltaSILAC measurements on more samples and relevant models is thus appealing in the future.

As for potential applications, DeltaSILAC could be quickly adapted to study protein modifications other than phosphorylation. However, the pSILAC design will limit this approach in cell line models or other systems where the isotopic labeling is possible^45, 46^. Also, mechanistically, more time points of pSILAC labeling and subsequent biochemical experiments are required to further understand more detailed or alternative mechanisms underlying the altered turnover rates. Examples of interesting questions are. e.g., whether the newly synthesized proteins are less likely to be phosphorylated or modified? Do modified and non-modified proteins follow different exponential decay kinetics^47^? How does the DeltaSILAC result change between different subcellular compartments?

In conclusion, DeltaSILAC offers a new and high potential workflow for timing the endurance of thousands of modified proteins, providing complementary knowledge to existing measurements for understanding functions of protein PTMs.

## Methods

### Cell culture

Twelve of 14 HeLa strains used in the multi-omics study were used in this report^26^, analyzing phosphoproteomic variability. The collection includes the original six HeLa cells variants subtype CCL2 (2, 6, 7, 12, 13, and 14) and six HeLa cells variants subtype Kyoto (1, 3, 4, 8, 9, and 10). The original HeLa_11 (deviating genome^26^) and HeLa_5 (CCL2.2) were not used in this study. All HeLa cells were gifts from multiple laboratories, as documented previously. The human colon cancer cell lines, SW948 and RKO, were kindly provided by Dr. Prem Subramaniam at Columbia University. All cells were tested and confirmed to be negative for mycoplasma. All cells were cultured for up to five additional passages for aliquoting and protein harvest. The cell culture protocol was detailed previously^26^. In brief, cells were routinely cultured in 5% CO_2_, and 37° in DMEM Medium supplemented with 10% FBS, Sigma Aldrich, together with a penicillin/streptomycin solution (Gibco).

### Pulsed SILAC experiment

For the two HeLa cell lines (HeLa_7, HeLa_8), SILAC DMEM medium (Thermo #88364) lacking L-arginine and L-lysine was firstly supplemented with 10% dialyzed FBS (Thermo Fisher, # 26400044) and the same penicillin/streptomycin mix. For both SW948 and RKO cell lines, the SILAC RPMI-1640 media lacking L-Arginine, L-Lysine (Thermo Scientific, # 88365) was used instead, with the same basic configuration. The Heavy L-Arginine-HCl (13C6, 15N4, purity >98%, #CCN250P1), and L-Lysine-2HCl (13C6, 15N2, purity >98%, # CCN1800P1) were purchased from Cortecnet and spiked into the culturing medium in the same manner as described previously for DMEM or RPMI medium^26, 48^. Before SILAC labeling, cells were seeded (at 40-50% confluency for CRC cells and 50-60% for HeLa cells) and incubated in normal light DMEM medium for 24 hours, at 5% CO_2_ 37°C, overnight. For pSILAC labeling, five-time points, including 0 hours and four labeling points 1, 4, 8, and 12 hours were applied, with three biological replicates (as individual dishes) per time point per cell line. This means a total of fifteen 10-cm dishes per cell line were prepared before labeling. For all four cancer cells, each replicate sample per time point yielded corresponding protein amount, sufficiently supporting 700 μg of protein mixture (see below) processed for proteomic and phosphoproteomic analysis.

### DeltaSILAC proteomic sample preparation

Label-free and labeled cells of HeLa cells, including HeLa_7 and HeLa_8, SW948, RKO, were harvested and digested, mainly as previously described^48, 49^. Cells were washed three times by precooled PBS, harvested, and snap-frozen by liquid nitrogen. The cell pellets were immediately lysed by adding 10 M urea containing complete protease inhibitor cocktail (Roche) and Halt™ Phosphatase Inhibitor (Thermo) and stored in −80°C for further analysis. After harvesting all dishes, samples of HeLa_7 and HeLa_8, as well as samples from SW948 and RKO, were processed for tryptic digestion. The cell pellets were ultrasonically lysed by sonication at 4 °C for 2 min using a VialTweeter device (Hielscher-Ultrasound Technology)^25, 48, 49^ and then centrifuged at 18,000 × g for 1 hour to remove the insoluble material. A total of 700 μg supernatant proteins (determined by BioRad Bradford assay) were transferred to clean Eppendorf tubes. The supernatant protein mixtures were reduced by 10 mM tris-(2-carboxyethyl)-phosphine (TCEP) for 1 hour at 37 °C and 20 mM iodoacetamide (IAA) in the dark for 45 min at room temperature Then five volumes of precooled precipitation solution containing 50% acetone, 50% ethanol, and 0.1% acetic acid were added to the protein mixture and kept at − 20 °C overnight. The mixture was centrifuged at 18,000×g for 40 min. The precipitated proteins were washed with 100% acetone and 70% ethanol with centrifugation at 18,000×g, 4°C for 40 min, respectively. 300 μL of 100 mM NH_4_HCO_3_ was added to all samples, which were digested with sequencing grade porcine trypsin (Promega) at a ratio of 1:20 overnight at 37 °C. After digestion, the peptide mixture was acidified with formic acid and then desalted with a C18 column (MarocoSpin Columns, NEST Group INC). The amount of the final peptides was determined by Nanodrop (Thermo Scientific). About 5% of the total peptide digests were kept for total proteomic analysis.

### DeltaSILAC phosphoproteomic sample preparation

From the same peptide digest above, 95% of peptides per sample was used for phosphoproteomic analysis. The phosphopeptide enrichment was performed using the High-Select™ Fe-NTA kit (Thermo Scientific, A32992) according to the kit instructions, as described previously^50^. Briefly, the resins of one spin column in the kit were divided into five equal aliquots, each used for one sample. The peptide-resin mixture was incubated for 30 min at room temperature and then transferred into the filter tip (TF-20-L-R-S, Axygen). The supernatant was removed after centrifugation. Then the resins adsorbed with phosphopeptides were washed sequentially with 200 µL × 3 washing buffer (80% ACN, 0.1% TFA) and 200 µL × 3 H_2_O to remove nonspecifically adsorbed peptides. The phosphopeptides were eluted off the resins by 100 µL × 2 elution buffer (50% ACN, 5% NH_3_•H_2_O). All centrifugation steps above were conducted at 500 g, 30sec. The eluates were collected for speed-vac and dried for mass spectrometry analysis.

### DIA mass spectrometry

For each proteomic and phosphoproteomic sample generated by DeltaSILAC, DIA-MS analysis was performed on 1 μg of peptides, as described previously^30, 48^.

LC separation was performed on EASY-nLC 1200 systems (Thermo Scientific, San Jose, CA) using a self-packed analytical PicoFrit column (New Objective, Woburn, MA, USA) (75 µm × 50 cm length) using C18 material of ReproSil-Pur 120A C18-Q 1.9 µm (Dr. Maisch GmbH, Ammerbuch, Germany). A 120-min measurement with buffer B (80% acetonitrile containing 0.1% formic acid) from 5% to 37% and corresponding buffer A (0.1% formic acid in H_2_O) during the gradient was used to elute peptides from the LC. The flow rate was kept at 300 nL/min with the temperature-controlled at 60 °C using a column oven (PRSO-V1, Sonation GmbH, Biberach, Germany).

The Orbitrap Fusion Lumos Tribrid mass spectrometer (Thermo Scientific) instrument coupled to a nanoelectrospray ion source (NanoFlex, Thermo Scientific) was calibrated using Tune (version 3.0) instrument control software. Spray voltage was set to 2,000 V and heating capillary temperature at 275 °C. All the DIA-MS methods consisted of one MS1 scan and 40 MS2 scans of variable isolated windows. This schema is comprised of 350∼373, 372∼391, 390∼407, 406∼421, 420∼434, 433∼447, 446∼459, 458∼470, 469∼482, 481∼494, 493∼506, 505∼517, 516∼529, 528∼541, 540∼553, 552∼564, 563∼576, 575∼588, 587∼602, 601∼614, 613∼626, 625∼641, 640∼655, 654∼669, 668∼683, 682∼698, 697∼714, 713∼732, 731∼751, 750∼771, 770∼792, 791∼815, 814∼839, 838∼868, 867∼899, 898∼939, 938∼983, 982∼1044,1043∼1127, 1126∼1650 m/z, with 1 m/z overlapping between windows. The MS1 scan range is 350 – 1650 m/z, and the MS1 resolution is 120,000 at m/z 200. The MS1 full scan AGC target value was set to be 2.0E5, and the maximum injection time was 100 ms. The MS2 resolution was set to 30,000 at m/z 200 with the MS2 scan range 200 – 1800 m/z, and the normalized HCD collision energy was 28%. The MS2 AGC was set to be 5.0E5, and the maximum injection time was 50 ms. The default peptide charge state was set to 2. Both MS1 and MS2 spectra were recorded in profile mode.

### Mass spectrometry data analyses

DIA-MS data analyses were performed using Spectronaut v13^51, 52^. For both proteomic and phosphoproteomic measurements, the hybrid assay libraries were respectively generated, which were based on both DIA measurements as well as data-dependent acquisitions (DDA) on relevant samples method described previously^48^), as well as the optimized pSILAC-DIA workflow^25, 27^. To generate the library for pSILAC DIA-MS datasets, the default settings for Pulsar search of Spectronaut was used with modification in the Labeling setting: a) “Labelling Applied” option was enabled, b) SILAC labels (“Arg10” and “Lys8”) were specified in the second channel. c) The complete H/L labeling of the whole library was ensured by selecting the “In-Silico Generate Missing Channels” option in the Workflow settings. d) Importantly, for phosphoproteomic datasets, the possibility of Phosphorylation at S/T/Y was enabled during database searching (as a variable modification), together with Oxidation at methionine was set as variable modification, whereas carbamidomethylation at cysteine was set as a fixed modification. The final spectral libraries (HeLa proteome library containing 179,199 peptide precursor assays for 8,040 protein groups, HeLa phosphoproteome library containing 139, 117 peptide precursor assays for 7,758 protein groups, CRC proteome library containing 78,763 peptide precursor assays for 6,593 protein groups, and CRC phosphoproteome library containing 56,056 peptide precursor assays for 4,859 protein groups) were all made public through PRIDE (see below).

For the targeted data extraction and subsequent identification and quantification for pSILAC datasets, the Inverted Spike-In (ISW) workflow was used, as described previously^27^. This means the “Spike-In” workflow was selected in Multi-Channel Workflow Definition, and both “Inverted” and “Reference-based Identification” options were enabled. Both peptide and protein FDR cutoff (Qvalue) were controlled at 1%%, and the data matrix was strictly filtered by Qvalue. The “minor peptide group” was defined as Modified Sequences. In particular, the PTM localization option in Spectronaut v13 was enabled to locate phosphorylation sites^31, 53^, with the probability score cutoff >0.75^31^, resulting Class-I peptides^4^ to be identified and quantified. All the other settings in Spectronaut were kept as Default.

### pSILAC based calculation for endurance analysis

In a pSILAC experiment analyzing protein turnover, because the growing cells are respectively maintained in a steady state^29^, it is assumed that the degraded and synthesized protein copies are balanced^28^. Accordingly, almost all (if not all) the proteoforms, including alternative splicing isoforms^33^ and proteins with PTMs, should achieve the concentration balance between synthesis and degradation in such a state (i.e., without any perturbation). This principle essentially enables the protein turnover calculation by monitoring the intensities of light and heavy peptides across several time points.

To fit the model of protein turnover estimation, we used a similar approach as was employed in our previous studies^25, 26^. Below, we describe such an approach in the present study.

a. We quantified peptide precursor intensities for light and heavy signals from the above Spectronaut results.
b. We calculated the *rate of loss* of the light isotope (k_Loss_) by modeling the relative isotope abundance (RIA, analogous to Pratt et al. ^54^). RIA is determined by the signal intensity in the light channel divided by the sum of light and heavy intensities (***RIA = L/ (H+L)***), onto an exponential decay model assuming a null heavy intensity (RIA = 1) at time 0; i.e., ***RIA(t)=e***^**− *klosst***^.
c. We used nonlinear least-squares estimation to perform the fit. A weighted average of the peptide precursor k_Loss_ values was performed to calculate the k_Loss_ values for all unique peptide sequence precursors, which were then aggregated as below.
d. We then summarized k_Loss_ to evaluate the turnover rate for each bulk-protein as previously described ^25, 26^, but also for each peptide and especially phosphopeptide in this study. In particular, for k_Loss_ determination, we applied a strict filter strategy as previous studies ^33^ by only accepting those peptide H/L ratios showing monotone increasing pattern during pSILAC labeling process and by only accepting those turnover rates with the CV of log2(k_Loss_) ^29^ <20% across three biological replicates.
e. To calculate the half-life T_1/2_^33, 54, 55^ for each peptide and phosphopeptide, we used ***T***_***1/2***_***= Ln(2)/ k***_***Loss***_ so that the k_Loss_ rate can be converted to a time domain. It should be noted that DeltaSILAC determination is performed per cell line. Thus, the relative doubling time difference between cell lines does not impact the calculation of our T_1/2_ values, which are essentially identical to the previously reported T_50%_ values^33^.
f. Finally, to pinpoint the endurance influence of each PTM, we subtracted T_1/2_ of backbone sequence-matching non-phosphorylated peptide (**np-peptide**) from the T_1/2_ of counterpart phosphopeptide (**p-peptide**), resulting in the value of ΔT_1/2_. This means ***ΔT***_***1/2***_ ***= T***_***1/2 p-peptide***_ ***-T***_***1/2 np-peptide***_ was applied.

The whole process of calculating T_1/2_ is also illustrated in **Figure S2**.

### Bioinformatics analyses

Circos-0.69-9 (http://circos.ca/) was used for the circle visualization **(Fig 1e**). Spearman’s correlation coefficients were calculated using R (functions cor() or cor.test() to infer statistical significance). The colored scatterplots from blue-to-yellow were visualized by the “heatscatter” function in R package “LSD” using a two-dimensional Kernel Density Estimation (**Figs 1g** and **6a-c**). The correlation matrix was created by the “corrgram” function in R package “corrgram” (**Fig 1f**). Partial correlation was analyzed by the “pcor” function in R package “ppcor” to inspect relationship between two continuous variables whilst controlling for or correct against the effect of another continuous variable. The melting temperature (Tm, °C) value (**Fig 4a**) for each phosphosite was taken from the reported dataset^6^. The flanking amino acid sequences (±14 amino acids) of phosphorylation sites were retrieved by motifeR (https://www.omicsolution.org/wukong/motifeR/)^56^. The Calculate the grand average of hydropathy (GRAVY) value for protein sequences, defined by the sum of hydropathy values of all amino acids divided by the protein length, was computed by GRAVY Calculator (http://www.gravy-calculator.de/index.php). The flanking amino acid sequences (±14 amino acids) of phosphorylation sites that can be unambiguously assigned to specific serine, threonine, or tyrosine residues were used for GRAVY calculation (**Fig 4b**). Secondary structure element, solvent accessibility, and transmembrane topology were predicted using Predict_Property standalone package (v1.01)^57^ with the protein FASTA sequences (**Figs 4e-g**). The functional score can reflect the importance of phosphosite for organismal fitness^11^. The sift score predicts the functional impact of missense variants based on sequence homology and the physicochemical properties of the amino acids. Lower scores represent deleterious variants. For every phosphorylated protein site, the functional score (**Fig 5a**), sift scores of alanine mutant (**Fig 5b**), the average sift scores of all variants, protein domains (**Fig 4c**), and kinase substrate motifs (**Fig 5d**) were retrieved from the previous report^11^. Kinase family tree (**Fig 5c**) was depicted by Coral (http://phanstiel-lab.med.unc.edu/CORAL/)^58^. Functional annotation was carried out in David Functional Annotation Tool v6.8 (https://david.ncifcrf.gov/summary.jsp) with all detected proteins in this study as background (**Fig 5e**). The compound responses were enriched by phosphorylation site-specific functional enrichment through ssGSEA2.0/PTM-SEA^59^. Sequence analysis (**Figs 5f and 6d**) was conducted and visualized by IceLogo (https://iomics.ugent.be/icelogoserver/)^60^. The modifiable ubiquitination sites in the flanking sequence (±14 amino acids) of the phosphosite were identified according to the reported dataset^42^, and the percentage of modifiable ubiquitination sites was calculated as the ratio of the number of ubiquitination sites over the total number of Lys (**Fig 5h**). The cellular compartment location of proteins with phosphorylation was annotated by a subcellular map of the human proteome^41^. All boxplots were generated using the R package “ggplot2”. The bold line within box indicates median value; box borders represent the first and third quartile, and whiskers and grey panels represent the minimum and maximum value within 1.5 times of interquartile range. Outliers are depicted using hollow dots. The heatmap was created using the R package “heatmap.2”.

### Data availability

The mass spectrometry data, all the spectral libraries used, and raw output tables as results have been deposited to the ProteomeXchange Consortium via the PRIDE^61^ partner repository.

## Author contribution

C.W. performed cell-culturing work of all HeLa cells, conducted pSILAC experiments for all cancer cell lines used, and prepared all the proteomics samples, with help from E.G., who established the CRC cell cultures. W.L. performed the DIA-MS and other MS measurements. Q.B., W.L. B.S., P.H., and Y.L. analyzed the MS data and performed bioinformatics analysis. Q.B. and Y.L. lead the bioinformatic analysis. Q.B. prepared most illustrations for the figures. T.M., G.R., and Y.D. provided critical inputs to the study and manuscript. C.W., W.L., and Y.L conceptualized the study. Y.L. wrote the paper with the inputs from all the other authors. Y.L. supervised the study.

## Acknowledgments

We thank Mark A. Lemmon and Evan G. Williams for discussions and critical comments on the manuscript. We thank Lukas Reiter and Tejas Gandhi for technical support in Spectronaut software. Y.L. was supported by a pilot grant from Cancer Systems Biology@Yale (CaSB@Yale) and a pilot grant from Yale Cancer Center at Yale University. C.W. was supported by International Visiting Program for Excellent Young Scholars of Sichuan University, China. B.S. was supported by a grant of the ASCR, v. v. i. (L200521953).

## Supplementary Figure Legend

**Supplementary Figure 1.**
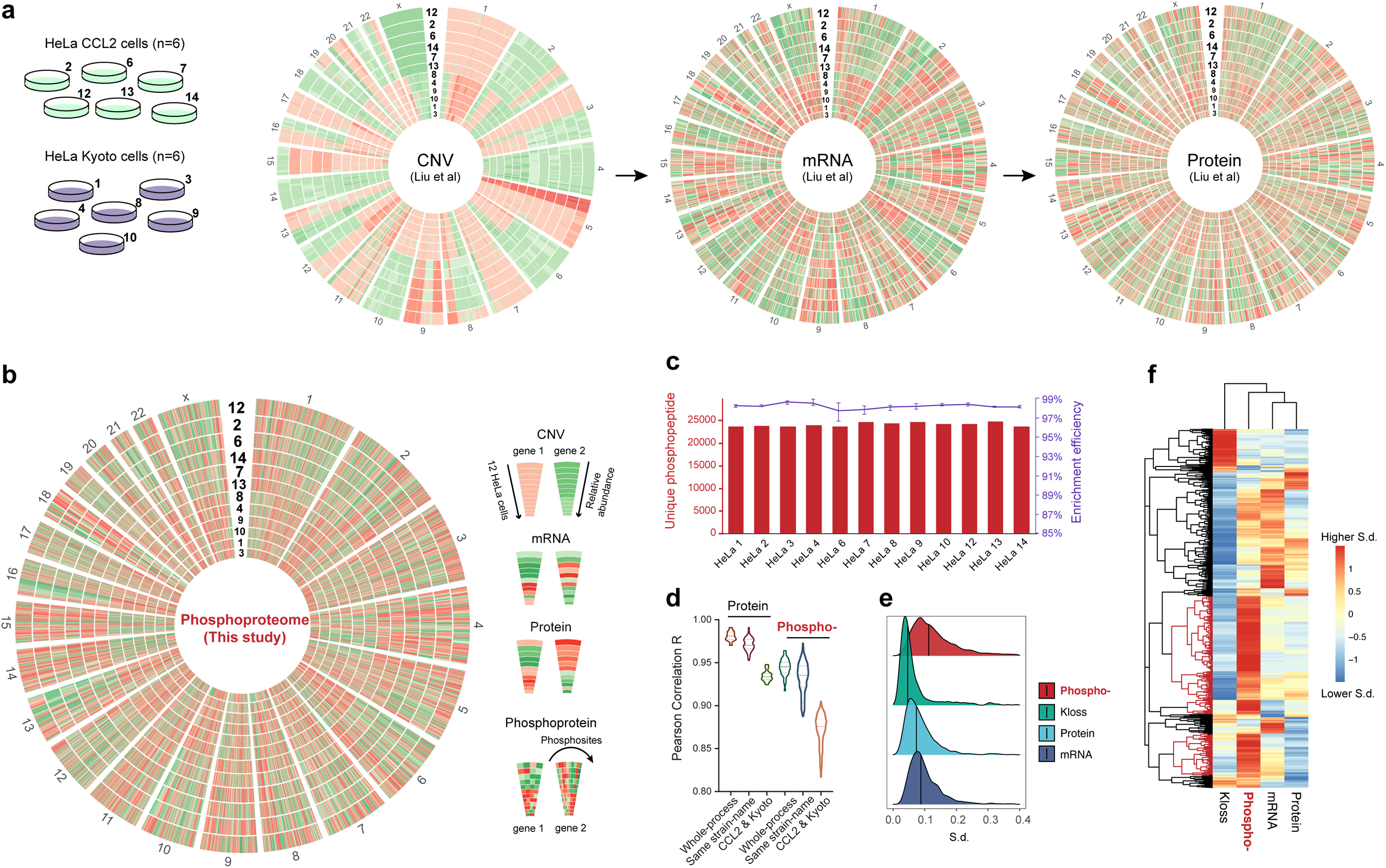
Phosphorylation profiles show high heterogeneity across 12 HeLa cell lines in steady states by the reliable DIA-MS measurement.

**Supplementary Figure 2.** Comparison of k_Loss_ and T_1/2_ profiles between HeLa_7 and 8 cells.

**Supplementary Figure 3.** Intra-protein variability and biological repeatability for ΔT_1/2_ values.

**Supplementary Figure 4.** Comparison of protein abundance and phosphoprotein endurance and melting temperature for site-specific phosphorylation.

**Supplementary Figure 5.** Endurance distribution quantified by DeltaSILAC in HeLa_7 and HeLa_8 cells.

**Supplementary Figure 6.** Distribution of ΔT_1/2_ values for phosphosites within different protein domains.

**Supplementary Figure 7.** Global relationships between local phosphosite environment and altered protein endurance.

**Supplementary Figure 8.** Distribution of conservation for phosphoamino acids with different endurance.

**Supplementary Figure 9.** Depiction of the kinome with different phosphoprotein endurance.

**Supplementary Figure 10.** Comparison of phosphoproteome endurance with biological features.

**Supplementary Figure 11.** Categories and functional analysis of phosphorylated protein endurance profiles.

**Supplementary Figure 12.** Distribution of ΔT_1/2_ values for phosphoproteins with different subcellular location.

**Supplementary Figure 13.** Phosphorylation increases endurance for most proteins in SW948 and RKO cells.

**Supplementary Figure 14.** Analysis of the connection between phosphoproteome endurance with compounds responses.

## References

1. Purvis, J.E. & Lahav, G. Encoding and decoding cellular information through signaling dynamics. Cell 152, 945–956 (2013).

2. Smith, L.M., Kelleher, N.L. & Consortium for Top Down, P. Proteoform: a single term describing protein complexity. Nature methods 10, 186–187 (2013).

3. Smith, L.M. & Kelleher, N.L. Proteoforms as the next proteomics currency. Science 359, 1106–1107 (2018).

4. Olsen, J.V. et al. Global, in vivo, and site-specific phosphorylation dynamics in signaling networks. Cell 127, 635–648 (2006).

5. Betts, M.J. et al. Systematic identification of phosphorylation-mediated protein interaction switches. PLoS computational biology 13, e1005462 (2017).

6. Huang, J.X. et al. High throughput discovery of functional protein modifications by Hotspot Thermal Profiling. Nature methods 16, 894–901 (2019).

7. Krahmer, N. et al. Organellar Proteomics and Phospho-Proteomics Reveal Subcellular Reorganization in Diet-Induced Hepatic Steatosis. Developmental cell 47, 205–221 e207 (2018).

8. Hunter, T. The age of crosstalk: phosphorylation, ubiquitination, and beyond. Molecular cell 28, 730–738 (2007).

9. Nguyen, L.K., Kolch, W. & Kholodenko, B.N. When ubiquitination meets phosphorylation: a systems biology perspective of EGFR/MAPK signalling. Cell Commun Signal 11, 52 (2013).

10. Lahiry, P., Torkamani, A., Schork, N.J. & Hegele, R.A. Kinase mutations in human disease: interpreting genotype-phenotype relationships. Nature reviews. Genetics 11, 60–74 (2010).

11. Ochoa, D. et al. The functional landscape of the human phosphoproteome. Nature biotechnology (2019).

12. Suizu, F. et al. The E3 ligase TTC3 facilitates ubiquitination and degradation of phosphorylated Akt. Developmental cell 17, 800–810 (2009).

13. Verma, R. et al. Phosphorylation of Sic1p by G1 Cdk required for its degradation and entry into S phase. Science 278, 455–460 (1997).

14. Skowyra, D., Craig, K.L., Tyers, M., Elledge, S.J. & Harper, J.W. F-box proteins are receptors that recruit phosphorylated substrates to the SCF ubiquitin-ligase complex. Cell 91, 209–219 (1997).

15. Mann, M. & Jensen, O.N. Proteomic analysis of post-translational modifications. Nature biotechnology 21, 255–261 (2003).

16. Aebersold, R. & Mann, M. Mass-spectrometric exploration of proteome structure and function. Nature 537, 347–355 (2016).

17. Swaney, D.L. et al. Global analysis of phosphorylation and ubiquitylation cross-talk in protein degradation. Nature methods 10, 676–682 (2013).

18. Gillet, L.C. et al. Targeted data extraction of the MS/MS spectra generated by data-independent acquisition: a new concept for consistent and accurate proteome analysis. Molecular & cellular proteomics : MCP 11, O111 016717 (2012).

19. Rosenberger, G. et al. Statistical control of peptide and protein error rates in large-scale targeted data-independent acquisition analyses. Nature methods 14, 921–927 (2017).

20. Meyer, J.G. et al. PIQED: automated identification and quantification of protein modifications from DIA-MS data. Nature methods 14, 646–647 (2017).

21. Ong, S.E. et al. Stable isotope labeling by amino acids in cell culture, SILAC, as a simple and accurate approach to expression proteomics. Molecular & cellular proteomics : MCP 1, 376–386 (2002).

22. Schwanhausser, B. et al. Global quantification of mammalian gene expression control. Nature 473, 337–342 (2011).

23. Jovanovic, M. et al. Immunogenetics. Dynamic profiling of the protein life cycle in response to pathogens. Science 347, 1259038 (2015).

24. Liu, Y., Beyer, A. & Aebersold, R. On the Dependency of Cellular Protein Levels on mRNA Abundance. Cell 165, 535–550 (2016).

25. Liu, Y. et al. Systematic proteome and proteostasis profiling in human Trisomy 21 fibroblast cells. Nature communications 8, 1212 (2017).

26. Liu, Y. et al. Multi-omic measurements of heterogeneity in HeLa cells across laboratories. Nature biotechnology 37, 314–322 (2019).

27. Barbora, S. et al. Isoform-resolved Correlation Analysis Between mRNA Abundance Regulation and Protein Level Degradation. Molecular systems biology In Press (2020).

28. Welle, K.A. et al. Time-resolved Analysis of Proteome Dynamics by Tandem Mass Tags and Stable Isotope Labeling in Cell Culture (TMT-SILAC) Hyperplexing. Molecular & cellular proteomics : MCP 15, 3551–3563 (2016).

29. Claydon, A.J. & Beynon, R. Proteome dynamics: revisiting turnover with a global perspective. Molecular & cellular proteomics : MCP 11, 1551–1565 (2012).

30. Mehnert, M., Li, W., Wu, C., Salovska, B. & Liu, Y. Combining Rapid Data Independent Acquisition and CRISPR Gene Deletion for Studying Potential Protein Functions: A Case of HMGN1. Proteomics, e1800438 (2019).

31. Bekker-Jensen, D.B. et al. Rapid and site-specific deep phosphoproteome profiling by data-independent acquisition without the need for spectral libraries. Nature communications 11, 787 (2020).

32. Yang, P. et al. Multi-omic Profiling Reveals Dynamics of the Phased Progression of Pluripotency. Cell Syst 8, 427–445 e410 (2019).

33. Zecha, J. et al. Peptide Level Turnover Measurements Enable the Study of Proteoform Dynamics. Molecular & cellular proteomics : MCP 17, 974–992 (2018).

34. Cox, J. & Mann, M. MaxQuant enables high peptide identification rates, individualized p.p.b.-range mass accuracies and proteome-wide protein quantification. Nature biotechnology 26, 1367–1372 (2008).

35. Lim, M.Y., Paulo, J.A. & Gygi, S.P. Evaluating False Transfer Rates from the Match-between-Runs Algorithm with a Two-Proteome Model. Journal of proteome research 18, 4020–4026 (2019).

36. Liu, Y. et al. Multi-omic measurements of heterogeneity in HeLa cells across laboratories. Nature biotechnology (2019).

37. Bubb, M.R., Lenox, R.H. & Edison, A.S. Phosphorylation-dependent conformational changes induce a switch in the actin-binding function of MARCKS. J Biol Chem 274, 36472–36478 (1999).

38. Schluter, C. et al. The cell proliferation-associated antigen of antibody Ki-67: a very large, ubiquitous nuclear protein with numerous repeated elements, representing a new kind of cell cycle-maintaining proteins. J Cell Biol 123, 513–522 (1993).

39. Holt, L.J. Regulatory modules: Coupling protein stability to phopshoregulation during cell division. FEBS Lett 586, 2773–2777 (2012).

40. Ng, P.C. & Henikoff, S. SIFT: Predicting amino acid changes that affect protein function. Nucleic Acids Res 31, 3812–3814 (2003).

41. Thul, P.J. et al. A subcellular map of the human proteome. Science 356 (2017).

42. van der Wal, L. et al. Improvement of ubiquitylation site detection by Orbitrap mass spectrometry. Journal of proteomics 172, 49–56 (2018).

43. Martin-Perez, M. & Villen, J. Determinants and Regulation of Protein Turnover in Yeast. Cell Syst 5, 283–294 e285 (2017).

44. Toyama, B.H. et al. Identification of long-lived proteins reveals exceptional stability of essential cellular structures. Cell 154, 971–982 (2013).

45. Kruger, M. et al. SILAC mouse for quantitative proteomics uncovers kindlin-3 as an essential factor for red blood cell function. Cell 134, 353–364 (2008).

46. Hidalgo San Jose, L. & Signer, R.A.J. Cell-type-specific quantification of protein synthesis in vivo. Nature protocols 14, 441–460 (2019).

47. McShane, E. et al. Kinetic Analysis of Protein Stability Reveals Age-Dependent Degradation. Cell 167, 803–815 e821 (2016).

48. Li, W. et al. Assessing the Relationship Between Mass Window Width and Retention Time Scheduling on Protein Coverage for Data-Independent Acquisition. J Am Soc Mass Spectrom (2019).

49. Collins, B.C. et al. Multi-laboratory assessment of reproducibility, qualitative and quantitative performance of SWATH-mass spectrometry. Nature communications 8, 291 (2017).

50. Gao, Q. et al. Integrated Proteogenomic Characterization of HBV-Related Hepatocellular Carcinoma. Cell 179, 561–577 e522 (2019).

51. Bruderer, R. et al. Extending the limits of quantitative proteome profiling with data-independent acquisition and application to acetaminophen-treated three-dimensional liver microtissues. Molecular & cellular proteomics : MCP 14, 1400–1410 (2015).

52. Bruderer, R. et al. Optimization of Experimental Parameters in Data-Independent Mass Spectrometry Significantly Increases Depth and Reproducibility of Results. Molecular & cellular proteomics : MCP 16, 2296–2309 (2017).

53. Rosenberger, G. et al. Inference and quantification of peptidoforms in large sample cohorts by SWATH-MS. Nature biotechnology 35, 781–788 (2017).

54. Pratt, J.M. et al. Dynamics of protein turnover, a missing dimension in proteomics. Molecular & cellular proteomics : MCP 1, 579–591 (2002).

55. Rost, H.L. et al. TRIC: an automated alignment strategy for reproducible protein quantification in targeted proteomics. Nature methods (2016).

56. Wang, S. et al. motifeR: An Integrated Web Software for Identification and Visualization of Protein Posttranslational Modification Motifs. Proteomics 19, e1900245 (2019).

57. Wang, S., Li, W., Liu, S. & Xu, J. RaptorX-Property: a web server for protein structure property prediction. Nucleic acids research 44, W430–435 (2016).

58. Metz, K.S. et al. Coral: Clear and Customizable Visualization of Human Kinome Data. Cell systems 7, 347–350 e341 (2018).

59. Krug, K. et al. A Curated Resource for Phosphosite-specific Signature Analysis. Molecular & cellular proteomics : MCP 18, 576–593 (2019).

60. Colaert, N., Helsens, K., Martens, L., Vandekerckhove, J. & Gevaert, K. Improved visualization of protein consensus sequences by iceLogo. Nature methods 6, 786–787 (2009).

61. Perez-Riverol, Y. et al. The PRIDE database and related tools and resources in 2019: improving support for quantification data. Nucleic Acids Res 47, D442–D450 (2019).

